# A workflow for visualizing human cancer biopsies using large-format electron microscopy

**DOI:** 10.1101/675371

**Authors:** Jessica L. Riesterer, Claudia S. López, Erin S. Stempinski, Melissa Williams, Kevin Loftis, Kevin Stoltz, Guillaume Thibault, Christian Lanicault, Todd Williams, Joe W. Gray

**Affiliations:** OHSU Center for Spatial Systems Biomedicine, Department of Pathology, Oregon Health and Sciences University, Portland, OR 97201

**Keywords:** Cancer, electron microscopy, FIB-SEM, SBF-SEM, 3DEM, volume rendering, tissue processing, large area mapping

## Abstract

Recent developments in large format electron microscopy have enabled generation of images that provide detailed ultrastructural information on normal and diseased cells and tissues. Analyses of these images increase our understanding of cellular organization and interactions and disease-related changes therein. In this manuscript, we describe a workflow for two-dimensional (2D) and three-dimensional (3D) imaging, including both optical and scanning electron microscopy (SEM) methods, that allow pathologists and cancer biology researchers to identify areas of interest from human cancer biopsies. The protocols and mounting strategies described in this workflow are compatible with 2D large format EM mapping, 3D focused ion beam-SEM and serial block face-SEM. The flexibility to use diverse imaging technologies available at most academic institutions makes this workflow useful and applicable for most life science samples. Volumetric analysis of the biopsies studied here revealed morphological, organizational and ultrastructural aspects of the tumor cells and surrounding environment that cannot be revealed by conventional 2D EM imaging. Our results indicate that although 2D EM is still an important tool in many areas of diagnostic pathology, 3D images of ultrastructural relationships between both normal and cancerous cells, in combination with their extracellular matrix, enables cancer researchers and pathologists to better understand the progression of the disease and identify potential therapeutic targets.

## Introduction

Electron microscopy (EM) is currently the only imaging technique that allows researchers to view the entire cellular ultrastructure at once with nanometer-resolution. Traditionally, transmission electron microscopy (TEM) has long been used as a pathology tool and method for viewing cellular ultrastructure (de Haro & Furness, 2012; Friedmann, 1961). However, TEM is a technique that is severely restricted in area and volume when standard 2D imaging, array tomography or conventional tomography is used (Han et al., 2017), and does not provide a complete picture of structures of interest. Cells, tissues, tumors, and organisms are all three-dimensional, and require 3D viewing to fully appreciate interactions and patterns in organization at the nanoscale (Association, 2019). Two scanning electron microscopy-based techniques are capable of collecting meaningful 3D volumes in a reasonable amount of time: focused ion beam-scanning electron microscopy (FIB-SEM) (Giannuzzi, 2005) and serial block face-scanning electron microscopy (SBF-SEM) (Denk & Horstmann, 2004).

3DEM provides an unparalleled view of tumor cell-cell interactions (Heymann et al., 2009), cell-extracellular matrix (ECM) interactions (Randles et al., 2016; Starborg et al., 2013) nuclear structure (Derenzini, Montanaro, & Treré, 2009), and organelle organization (Johnston et al., 2018; Novotný et al., 2013). However, three-dimensional electron microscopy (3DEM) has only found a footing in life sciences in the last decade, and is still not routinely used (Peddie & Collinson, 2014). Commercially available 3DEM instruments have finally been developed with enough control to image soft, beam-sensitive life science samples easily with minimal-to-no beam damage (Ishitani, Hirose, & Tsuboi, 1995; Jorgens et al., 2017; López et al., 2017; Midgett, López, David, Maloyan, & Rugonyi, 2017; Rennie, Gahan, López, Thornburg, & Rugonyi, 2014). Recent developments in instrumentation and computing power allow for large data sets generated by 3DEM to be collected for statistical analysis and comparison of translational tissue samples (Boergens et al., 2017; Brent & Boucheron, 2018; Falk et al., 2018; Johnston et al., 2018; Liu, Jones, Seyedhosseini, & Tasdizen, 2014).

Three-dimensional imaging is established in clinical settings to understand spatial relationships (Taha & Hanbury, 2015). Radiographs and CT scans are vital when diagnosing and treating cancer patients, and can be utilized when implementing personalized treatment plans. Personalized medicine will be executed more readily in the future thanks to novel clinical trials currently in progress (Johnson et al., 2018). Continued personalized medicine research for cancer treatment requires an intimate understanding of tumor behavior and architecture. In addition to the conventional clinical imaging, 3DEM of clinical specimens can elucidate tumor behavior and be linked to gene profiles and proteomic signatures (Cho, Irianto, & Discher, 2017). When coupled with large format mapping of tumor tissue, quantification of cancer cell behavior at the nanoscale can be achieved while also providing meaningful information regarding tumor heterogeneity at the macroscale.

To assist in this effort, a workflow is presented here that is straight-forward, utilizes common wet bench techniques, and produces high-resolution large-format datasets in 2- and 3-dimensions. While this manuscript focuses on cancer tumor specimens, the workflow can be applied to cell culture, xenografts, organoids, and other tissues from both human and animal models.

## Materials and Methods

The following workflow can be fully implemented, from tissue collection to final image collection (quantification excluded) in roughly two weeks. Figure 1 presents a schematic outline of this workflow. Proper fixation of clinical specimens is described and a reliable staining protocol is provided. Final geometries for trimmed resin-embedded specimen blocks are described with tips and tricks. Screening and review methods are summarized to save the researcher time and effort. Finally, electron microscopy imaging methodology is explained to achieve large-format high-resolution 2D maps and 3DEM.

**Figure 1:**
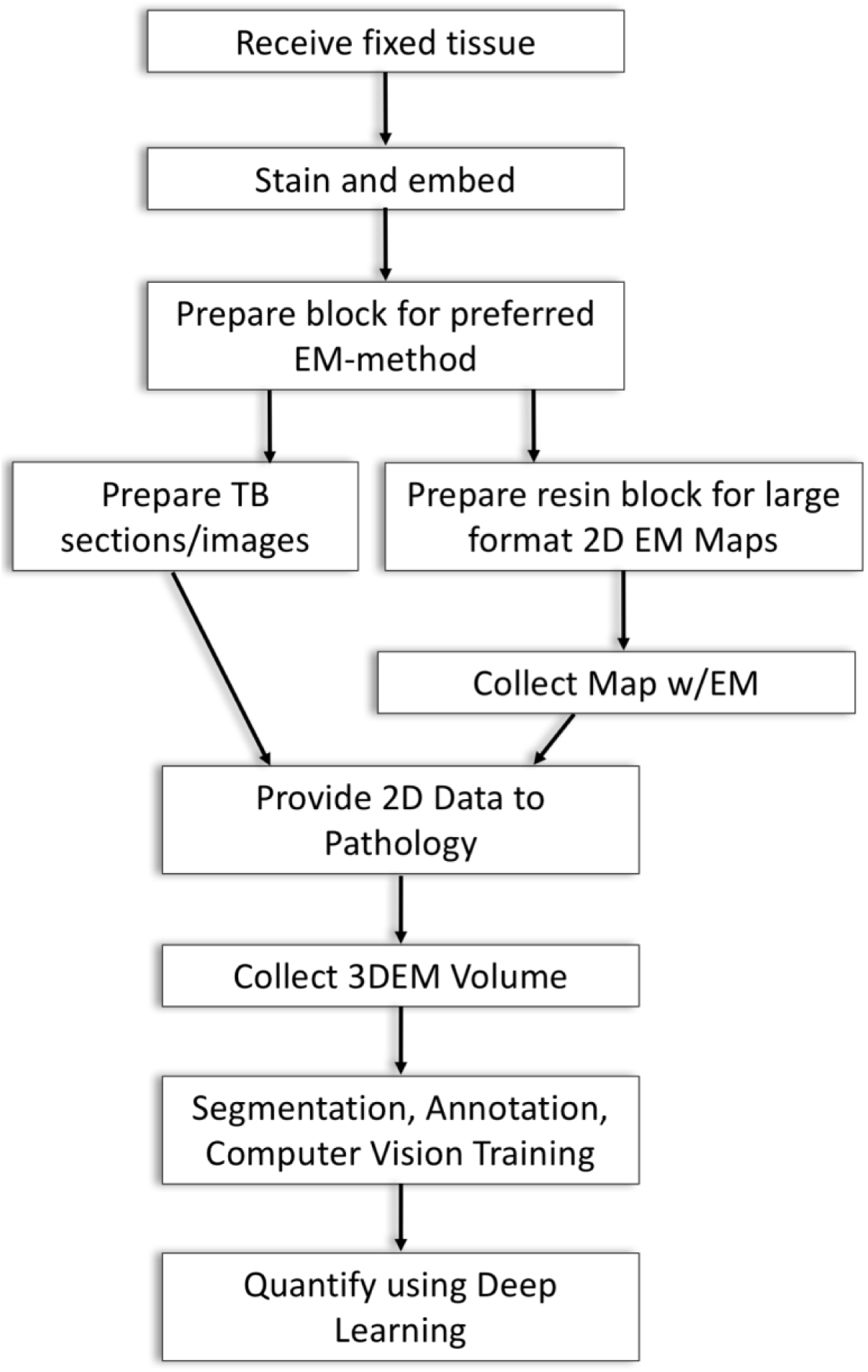
The 3DEM workflow utilizes standard techniques employed in wet bench processing and scanning electron microscopy. Tissue specimens, cell cultures, and organoids can all be imaged using the approach outlined. Fixation is the most critical step, and specimens must be placed in the fixative solution as quickly as possible; preferably within a few minutes after specimen collection.

Tissues, cell monolayers, organoids and xenografts all image differently depending on the sample preparation method utilized. Moreover, the same tissue type from different species may also need different fixation solution and processing methods to have optimal contrast and charge mitigation when the same microscope is used (Borrett & Hughes, 2016; Kizilyaprak, Longo, Daraspe, & Humbel, 2015; Kopek et al., 2017). For example, brain tissue may be processed successfully using a protocol that yields poor images when applied to cancer tissues (unpublished data). Researchers are therefore encouraged to dive into the literature and test new sample preparation protocols for a specific sample-type. Also, and if available, having the ability to evaluate the use of FIB-SEM versus SBF-SEM will help the researcher to design a data collection strategy. The protocols evaluated during development of this workflow included the Dresden protocol (Paridaen, Wilsch-Bräuninger, & Huttner, 2013), Renovo (Mukherjee et al., 2016) and the Hua method (Hua, Laserstein, & Helmstaedter, 2015). We settled on the the Hua method with some modifications as described below, for human cancer biopsies. The final protocol described is sufficient for large format mapping and 3DEM FIB-SEM and SBF-SEM, eliminating the need for multiple samples processed with different protocols. In the case of biopsy tissue, where sample acquisition is limited and precious, flexibility is an important advantage of this workflow.

### Sample Fixation

The most crucial step in the entire workflow is sample fixation. Tissue needs to be preserved in strong fixative as soon as possible to maintain cellular ultrastructure. In a clinical setting, fixation time is critical in order to capture precious human tissue adequately. Of course, priority is given to patient care, and therefore, the tissue sometimes cannot be handled as quickly as needed for optimal ultrastructure preservation. However, we recommend 2 minutes as a “best practice” time to start of preservation. In order to facilitate quick fixation, the team of clinical coordinators are provided with Eppendorf tubes containing 1.5 mL of fixative solution to have on-hand in the operating room during biopsies and resections. Karnovsky’s fixative (2.5% paraformaldehyde, 2.5% glutaraldehyde in 0.1M Na Cacodylate buffer (pH 7.4)) is the solution of choice (Karnovsky, 1965). Eppendorf tubes with and without tissue are stored at 4°C. Tissues collected here are typically 18-gauge core needle biopsies, where 3-4 mm of one core is preserved as described, as shown in Figure 2a.

**Figure 2:**
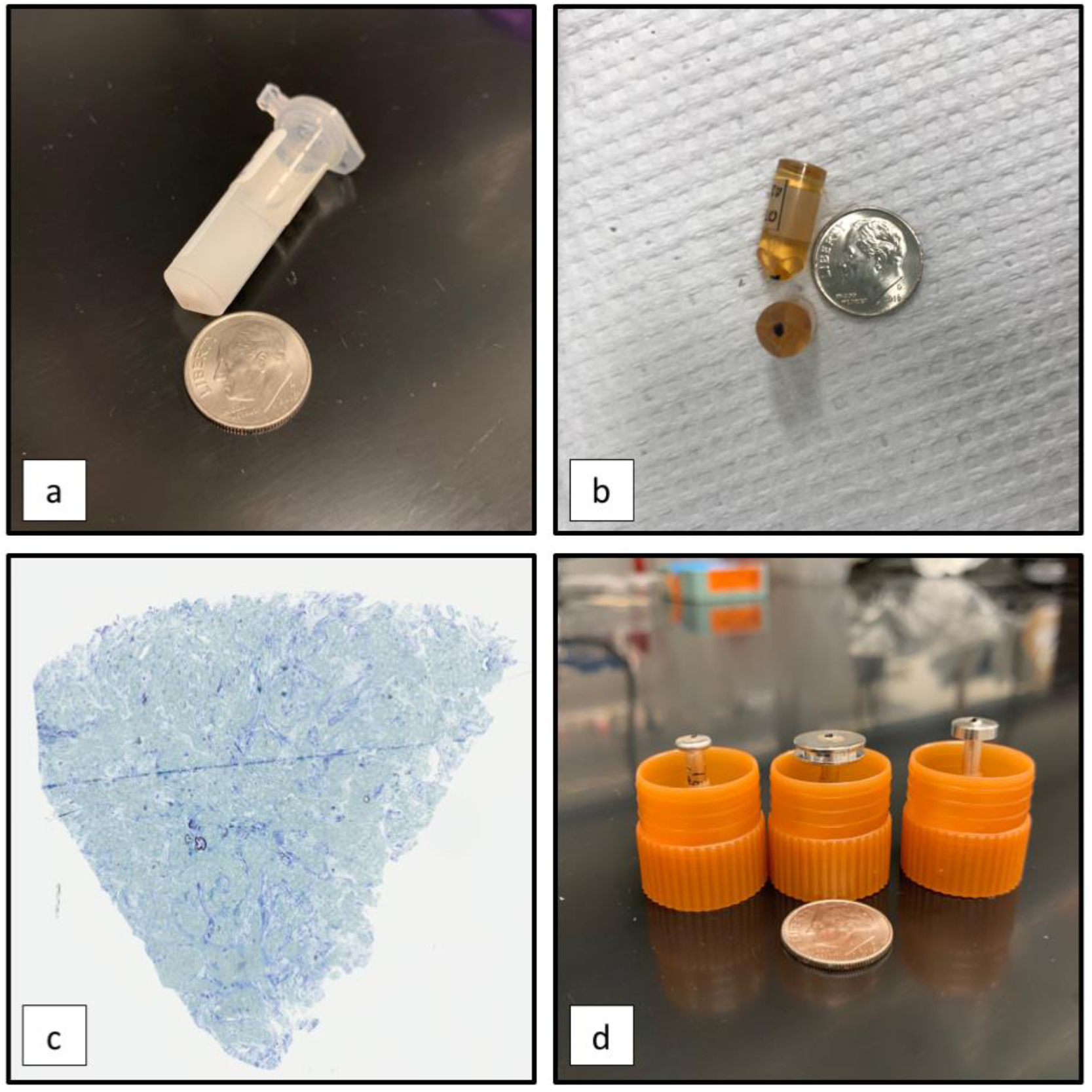
The sample takes several forms throughout the workflow. Only a **(a)** small piece (3-4 mm of 18-gauge core needle biopsy), as shown at the bottom of the fixative solution, is needed to produce several epoxy resin blocks **(b)**. Tissue will be black and opaque in the amber epoxy resin after the 3DEM protocol of choice is implemented. **(c)** TB sections for sample screening can be taken from samples mounted directly to **(d, l-r)** Mini Pin stubs or from traditionally molded blocks like in **(b)**. Mini Pin, standard 12 mm diameter, or Agar Microtome pin-style stubs may be used where only a small portion of the original tissue from **(a)** is mounted. Conductive coating of the entire sample and stub is the final step prior to imaging.

Typically, clinical tissues that need to be preserved quickly for protein assays or additional imaging via confocal techniques are flash frozen in the operating room (Fischer, Jacobson, Rose, & Zeller, 2008). Unfortunately, flash frozen specimens are not adequate for preserving cellular ultrastructure, as illustrated in Figure 3 (Scouten & Cunningham, 2006). Comparing flash frozen MCF7 cultured cells to those chemically fixed provides appreciation for fixation method. However, high pressure freezing (HPF) is a preferred fixation method and often gives superior ultrastructure preservation of cells, but is not feasible in a clinical setting (Shuo Li, Raychaudhuri, & Watanabe, 2017; McDonald, 2014).

**Figure 3:**
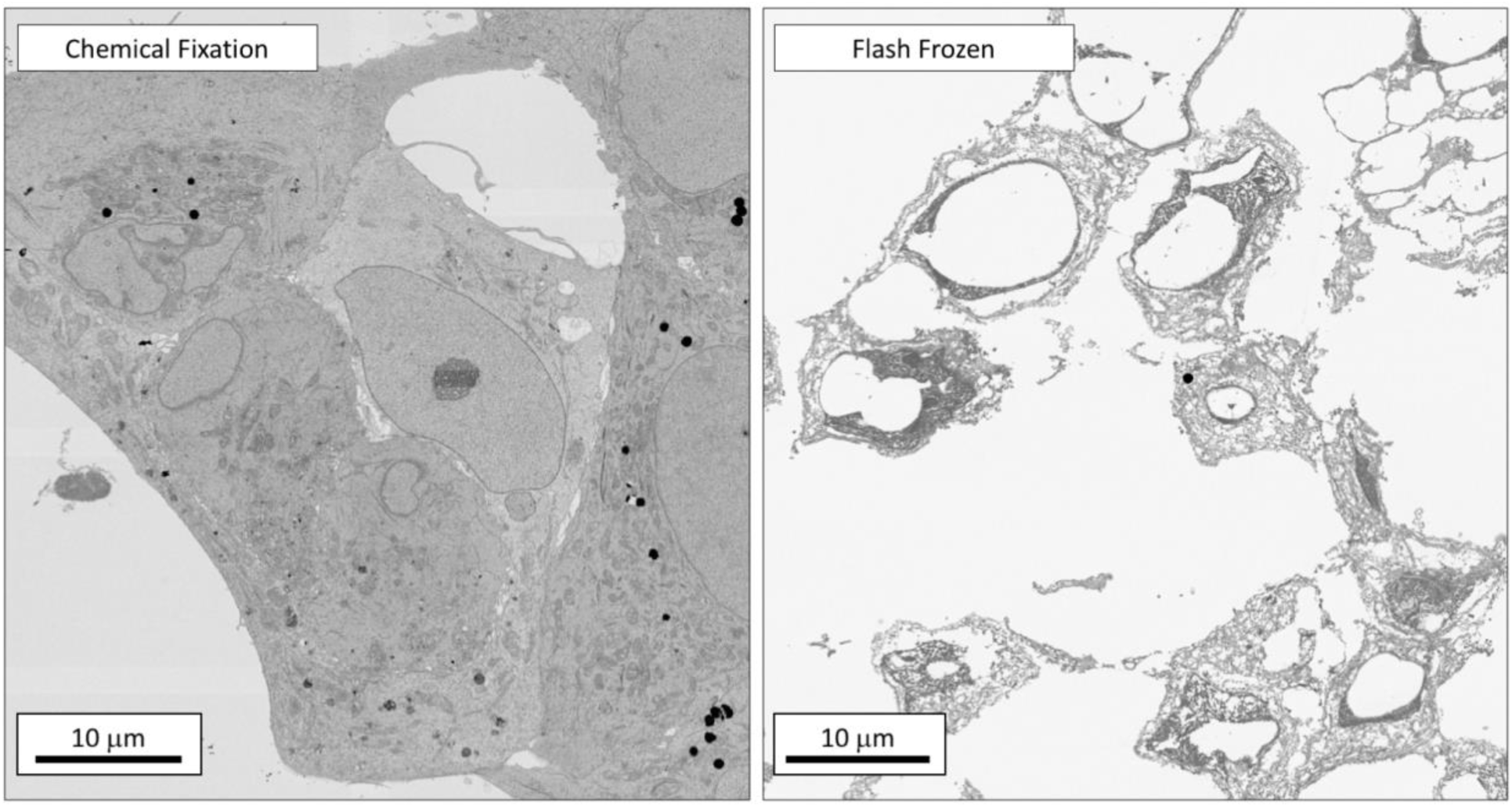
Proper fixation is crucial for meaningful data collection via 2D mapping or 3DEM. Chemical fixation maintains cellular ultrastructure and cell-cell relationships. While flash freezing is a standard technique to preserve information for other imaging and diagnostic techniques, the procedure is not fast enough to preserve regions of interest (ROIs) for EM. In addition, ice crystals formed during this freezing process can damage sub-cellular structures (Scouten & Cunningham, 2006).

### Post-fixation and Staining Protocol

#### Day 1

Once the fixed samples are received, they are post fixed in 2% (v/v) OsO_4_ prepared in 0.1M Na Cacodylate (pH 7.4) for 1.5 hours at room temperature. After this step, the samples are incubated for 1.5 hours at room temperature in 2.5% (w/v) potassium ferricyanide (K_3_[Fe(CN)_6_]) dissolved in 0.1M Na Cacodylate (pH 7.4). Samples are then extensively washed in dH_2_O for 5 minutes with 5 fresh exchanges. After washing, the samples are incubated in a conventional oven for 45 minutes at 40°C in freshly prepared 1% (w/v) thiocarbohydrazide (TCH) solution. This step is followed by another 5 exchanges of fresh dH_2_O over 25 minutes, and then incubated in 2% (v/v) OsO_4_ for 1.5 hours at room temperature. Water washes are then repeated as described above. At the end of day 1, the samples are incubated overnight in the dark at 4°C in 1% (w/v) aqueous uranyl acetate. All these steps are performed using aluminum foil protected 2 ml centrifuge tubes. The three incubations done at room temperature are completed using a rotating platform.

#### Day 2

Samples are moved from 4°C incubation into a conventional oven set up at 50°C for two more hours. After this, the samples are washed in dH_2_O for 5 minutes with 5 fresh exchanges. In the next staining step, the samples are transferred to a lead aspartate solution, previously warmed at 60°C, and incubated for 2 hours at 50°C using a conventional oven. Again, these steps are protected using aluminum foil covered 2 ml centrifuge tubes, as described above. Another 5 exchanges of fresh dH_2_O over 25 minutes are completed. For this washing step the initial rinse is done in dH_2_O warmed in a 60°C oven. The dehydration procedure is achieved in a series of acetone:water (50, 75, 85, 95%) mixtures with the final last step consisting of 100% acetone. It is worth mentioning that a new acetone bottle is utilized each time samples are processed using this protocol. Each dehydration step is completed twice for 5 minutes with one fresh exchange of the dehydrating solution. The samples are then infiltrated in epoxy resin (EMS EMbed 812 cat# 14120). For this, samples are incubated in 1:1 acetone:resin for 40 minutes using a rotating platform followed by a 1:3 mixture and the same incubation period. At the end of Day 2 the samples are transferred into vials with fresh 100% resin and incubated overnight on a rotating platform.

#### Day 3

Fresh epoxy resin is prepared and the samples are incubated with fresh resin for 30 minutes with 4 fresh exchanges. After resin infiltration, the 1 mm^2^ sample blocks are wicked onto filter paper (Whatman #1) to remove the excess resin (Schieber et al., 2017) and then placed onto “Mini Pin” gunshot residue stubs (Ted Pella, Inc. Product #16180). The samples are polymerized at 60°C for 48 hours using a conventional oven. If preferred, samples may also be preserved in a traditional BEEM capsule or mold and polymerized in the same manner.

#### Day 5

Samples are removed from the oven and allowed to cool to room temperature. The advantage of mounting the samples directly to the stub in this way is that there is no need for extra trimming to expose the sample block. Furthermore, these mini pins can be fitted to a microtome sample holder for trimming and sectioning as needed without risk to the trim tool or histology diamond knives. If traditional blocks from molds are prepared, the final result will be similar to those pictured in Figure 2b. If the researcher decided to polymerize the sample in a traditional BEEM capsule or mold, then the final sample needs to be trimmed and exposed as routinely done in an EM sample preparation laboratory.

### Toluidine blue (TB) screening

Once the samples have been processed for 3DEM the entire block face (∼1 mm^2^) is sectioned using a Leica UC7 ultramicrotome to generate 650 nm sections that are stained with toluidine blue solution (Sridharan & Shankar, 2012) and imaged using a conventional bright field optical microscope. This section has enough quality for the team of researchers and pathologists to decide if more sectioning is needed to expose areas of the sample that are deeper within the tissue, providing at the same time information regarding tissue quality and heterogeneity. Figure 2c shows an example. Since the TB section obtained using this method encompasses the entire block face, the acquired optical image can be easily overlaid with the EM map image, as describe below.

### SEM screening

After EM preparation, the samples described above can also be screened via traditional SEM imaging by mounting a semi-thin 250-350 nm section of the 1 mm^2^ block faces onto a glow discharged 5×7 mm silicon chip (Ted Pella, Inc., cat# 16007). For the silicon chip surface activation, we use an easiGlow PELCO™ unit at 15 mA for 30 seconds. Silicon chip screening has the same benefit as screening with TB, in that tissue quality and heterogeneity can be quickly assessed. Moreover, fixation and staining quality can be evaluated. Chips are mounted to standard 12 mm SEM stubs with carbon tape for SEM viewing (Ted Pella, Inc., cat# 16111 and 16084-7, respectively).

### Pathology Review

Consultation with a pathology team is a key step in the workflow, but finding a common language is a must. TB stained sections from the block face not only serve as a quick quality or location check, but also provides the link between a pathologist and the electron microscopist. Pathologists typically derive diagnostic information from images of hematoxylin and eosin (H&E) or immunohistochemistry (IHC) stained formalin fixed paraffin embedded (FFPE) tissue sections. However, these staining techniques cannot easily be applied to epoxy sections or tissue stained with heavy metals (Pool, 1969). TB staining is an adequate compromise.

Since tissue and extracellular features look much different in a color image at low-resolution compared to a high-resolution grayscale image, the two must be viewed together. As described in a following section, TB and EM images from adjacent sections can be overlaid onto one another for direct comparison. Software packages such as Thermo Scientific Maps™, Glencoe Software PathViewer™, and OMERO (Simon Li et al., 2016) allow easy digital image sharing and region of interest annotation.

### Sample trimming for large format mapping and 3DEM

Trimming the mounted block face for large format 2D mapping and 3DEM via FIB-SEM is straight-forward. As seen in Figure 2d, the block face needs to be as close to parallel to the SEM stub as possible. As mentioned previously, Mini Pin SEM stubs can be used on a standard microtome and allow adequate diamond knife clearance. Typically, a trim tool or histology knife is suitable for this and this step can be coupled with TB and SEM screening sections. In fact, the workflow works best when the screening sections are the last consecutive sections coming from the mounted block face. Additional trimming to tidy the block sides into a pillar can be performed, but is not necessary. NOTE: if standard 12 mm SEM stubs are being used, trimming and sectioning directly from a resin-embedded BEEM capsule or coffin mold formed block is advised prior to stub mounting, as the diamond knife will most likely collide with the larger diameter SEM stub.

Conductive coating is necessary to achieve high-resolution, high contrast, low noise images on samples that build up surface charge while imaged with an electron and/or ion beam. Carbon is preferred for FIB-SEM due to its amorphous nature, minimizing ion milling artifacts (i.e., curtaining) that distort final image quality.

Samples selected for SBF-SEM are mounted on Microtome stub SEM pins (Agar Scientific, cat# 61092450) using conductive silver epoxy (Ted Pella, Inc., H2OE EPO-TEK® cat# 16014) and trimmed using a Diatome trim90 diamond knife to generate a 500×500 µm^2^ pillar with the block face parallel to the stub. These SEM stubs are also small diameter, and can be used on the microtome similar to the Mini Pins. This sample is then coated with 20 nm of gold using the aforementioned sputter coater. Gold coating, especially on the exposed sides of the trimmed resin block, is preferred for SBF-SEM due to the superior electrical conductivity of metal over carbon (Nguyen et al., 2018).

A Leica ACE600 High Vacuum Sputter Coater fitted with a rotating platform was used in this study for carbon and gold coating. The coating thickness was monitored and measured with an *in situ* quartz crystal localized in the center of the rotating platform. Stubs are coated after trimming is complete.

### EM Imaging

Once a sample preparation protocol is established, imaging conditions for the respective 3DEM technique typically will not vary much for the same tissue or cell type. Depending on the exact microscope model, small variations will be needed from those reported here. While collecting a stunning image for analysis can be straightforward, image collection parameters must also be optimized for post-imaging data handling. Scanning conditions chosen here provide the best membrane contrast and signal-to-noise ratio, while preventing charging artifacts and being conscientious of data collection time. As recent image denoising experiments in the machine learning community show, computer vision algorithms struggle with low signal-to-noise and image-to-image contrast variations (Buades, Coll, & Morel, 2005; Remez, Litany, Gires, & Bronstein, 2017). Minimizing these variations at the start of data acquisition will make data segmentation and analysis easier.

3DEM can be collected using FIB-SEM and/or SBF-SEM. Example datasets, reconstructed using the respective techniques, are shown as screenshots in Figure 4. Movies showing the full capability of 3DEM are provided in the supplemental material for viewing. FIB-SEM provides roughly 25 × 20 × 8 µm^3^ volumes in 72 hours. Voxel sizes are isotropic at 4 nm resolution. These volumes are suitable for viewing cellular components like mitochondria, Golgi apparatus, and nucleoli in fine detail. In some cases, resolution may be sacrificed for more volume if larger microstructures, such as tumor nest organization with respect to ECM, are desired. In this case, SBF-SEM would be the preferred method. Volumes collected in 72 hours are roughly 60 × 40 × 30 µm^3^ with 10 nm lateral resolution and 40 nm depth resolution, i.e. slice thickness. If samples are initially prepared on Agar Microtome stubs as described above, both techniques may be utilized on the same specimen. Which technique to use is determined during TB and SEM screening and after consultation with the pathology consultant.

**Figure 4:**
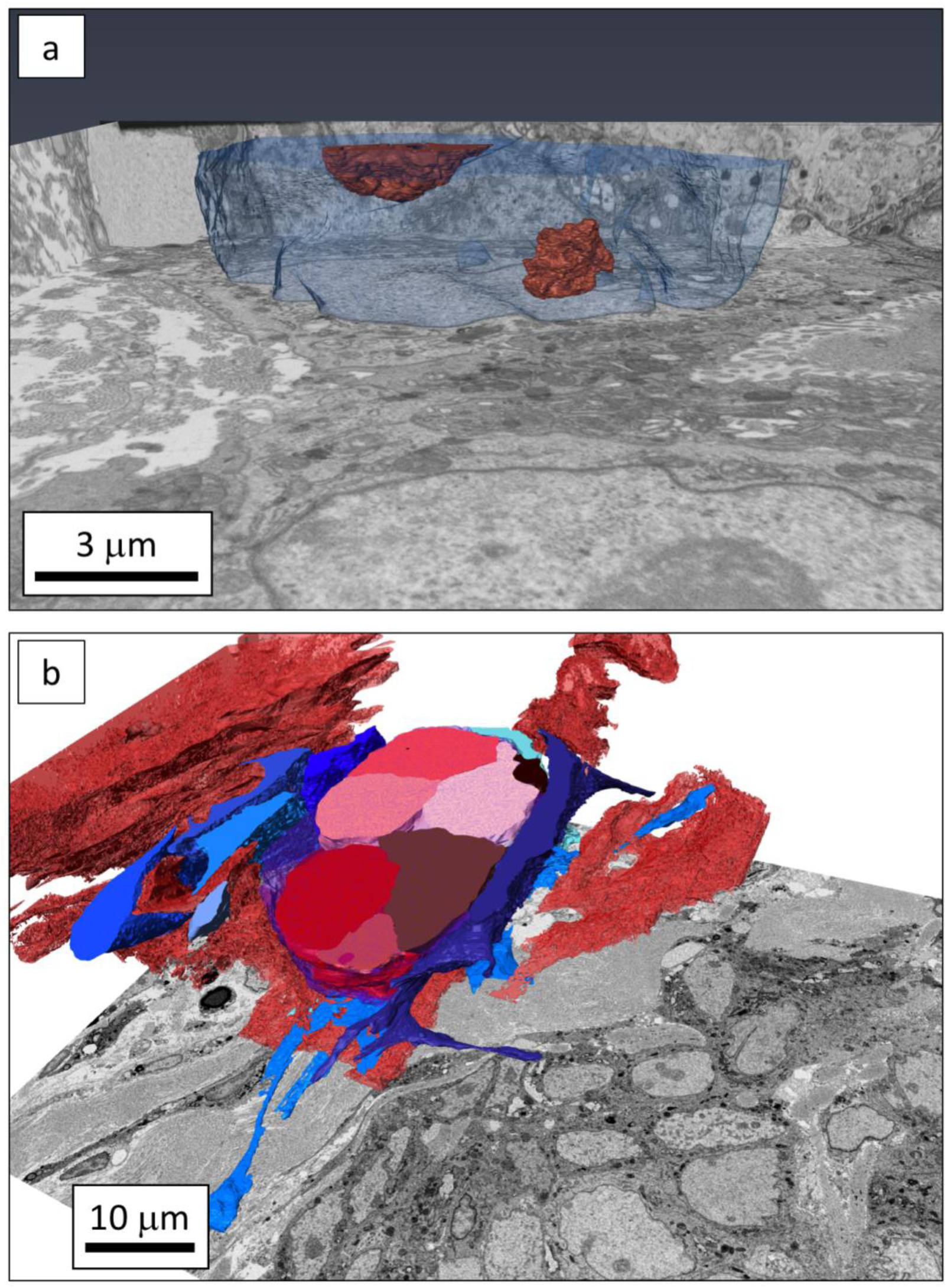
Once rendered, volumes generated from 3DEM datasets are rich in information. FIB-SEM data is presented in **a** and shows the fine detail of a cancer cell nucleus (blue) and nucleoli (orange). SBF-SEM datasets are larger volume, as shown in **b**. Here, the relationship between the microenvironment (red collagen fibers and blue fibroblasts) surrounding the center red and pink tumor cells is displayed.

### Large Format 2D Mapping

The Thermo Scientific Maps™ software for SEMs allows high resolution imaging (4 nm/pixel) across the entire block face by montaging hundreds of images together during automated image tile acquisition, as described in Figure 5. The resulting high-resolution map has panning and zooming capabilities and can be used like “Google Street-View” for the SEM. Stage positions are saved within the project, allowing specific ROIs identified earlier to be found with ease even if the sample has had to be moved from the microscope in the meantime. Images can be exported at full resolution, saved via screen captures, viewed as raw image tiles, and shared amongst collaborators using Microsoft HDView™ format.

**Figure 5:**
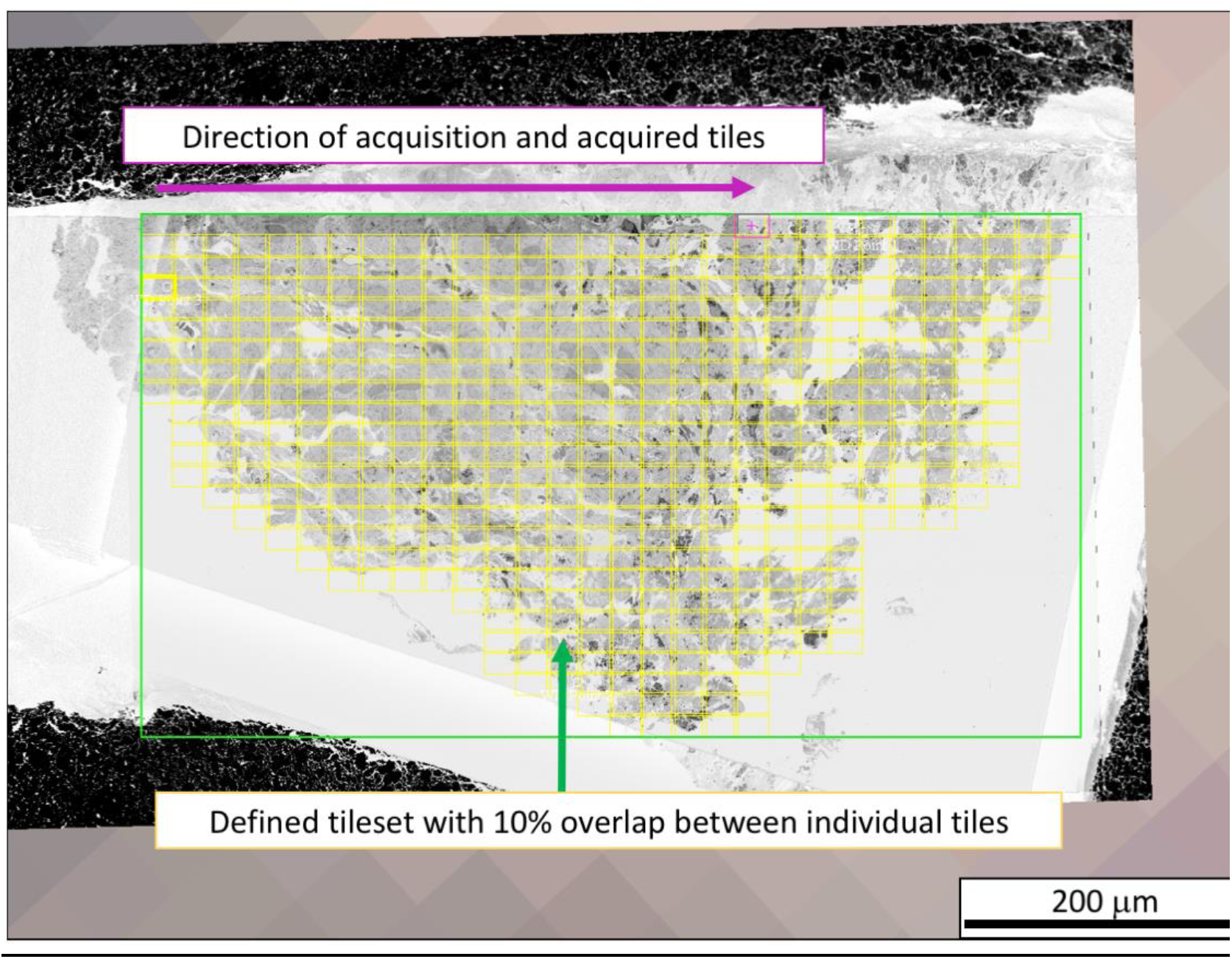
Large-format high-resolution 2D maps are collected over the entire block face at 4 nm/pixel spatial resolution. A tileset (yellow) of hundreds of images is defined over the sample. Individual tiles are automatically acquired and subsequently stitched together using the user-defined tile overlap. Within the Maps software, collected tilesets can be viewed by panning over and zooming into features of interest. Specific features can be marked and returned to for 3DEM data acquisition.

Large-format high-resolution maps are acquired over 15-24 hours depending on size of area being acquired (∼500 × 500 µm^2^). Typical imaging conditions used for data collection on an FEI Helios NanoLab 660 DualBeam™ FIB-SEM are 2-3 keV, 200-400 pA, 4 mm working distance with the retractable directional backscattered (DBS) electron detector. Tiling is done using 10% overlap in both *x* and *y* directions. Digitized TB section images are imported as tiff files and overlaid onto the EM montage via a Maps software-driven 3-point alignment procedure. The correlated images can be faded into each other by opacity adjustments, also providing aid in locating ROIs.

### High Resolution 3DEM via FIB-SEM

The FIB-SEM is also used to collect targeted 3D volumes using a heated Ga^+^ liquid metal ion source (LMIS) slicing 4 nm-thick for hundreds of slices (Giannuzzi, 2005). The slicing/imaging cycle is automated using the FEI AutoSlice and View™ software package. Figure 6 illustrates the experimental geometry. Samples are tilted such that the block face is perpendicular to the FIB (52° on a FEI/Thermo Scientific instrument) for thin slicing. Without moving the stage, a backscattered electron image is generated with the SEM after slicing is completed. The process repeats to form an image stack which is later aligned and annotated. Fiducial markers are placed near the ROI to aid in slice and image placement. A carbon or platinum protective capping layer is deposited over the volume to be collected in order to maintain the sample integrity during ion beam fiducial recognition. The largest data set for a single volume collected with in-house instrumentation is over 2900 images at 4 nm isotropic voxels, but FIB-SEM has been utilized by others for 3DEM under unique conditions for months at a time (Xu et al., 2017).

**Figure 6:**
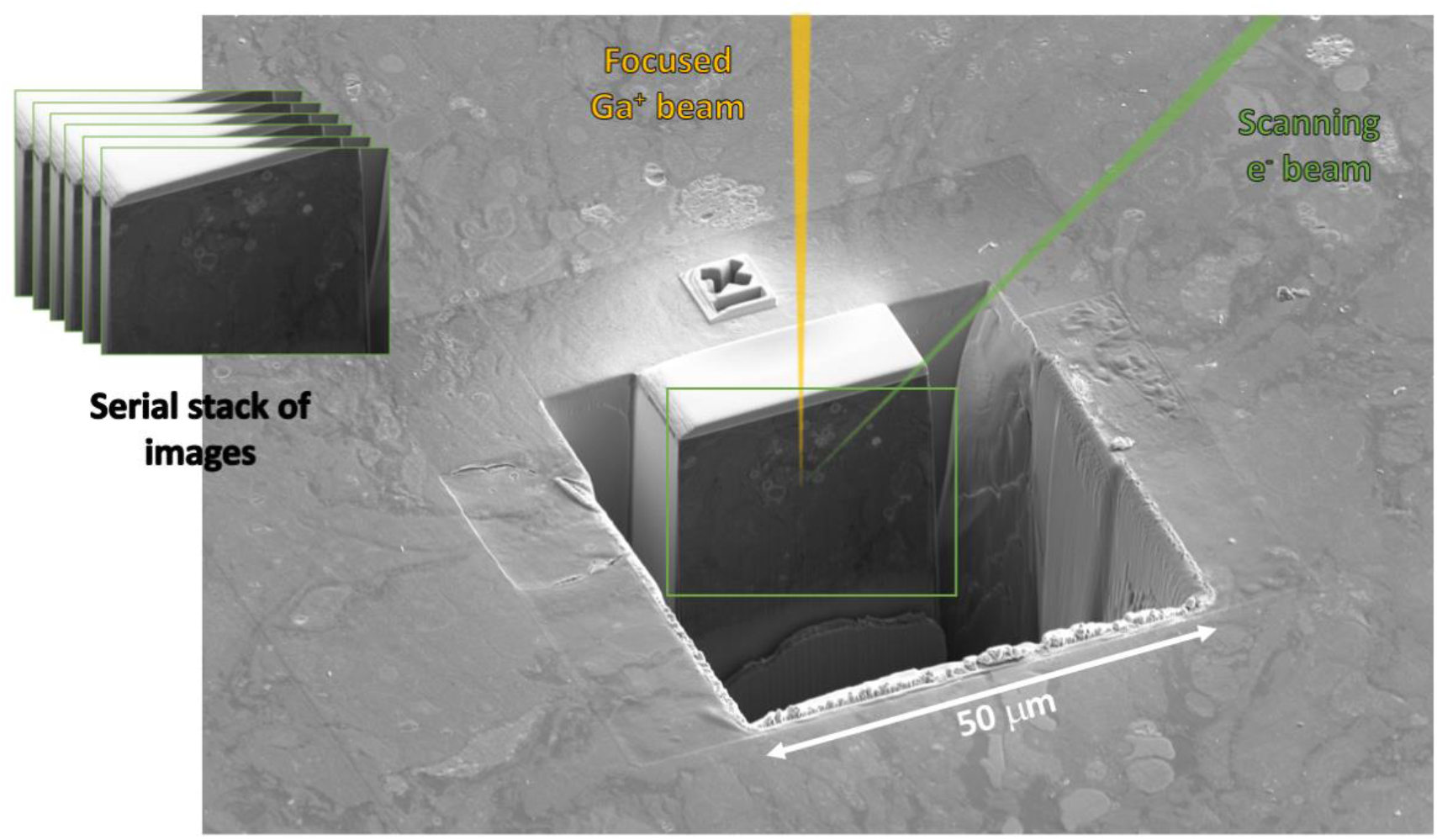
FIB-SEM utilizes coincident electron and Ga^+^ beams to serially section samples. The top surface of the block (previously mapped) is used for ROI identification. A protective capping layer (white box) is deposited over the volume to be collected, and a wide trench is milled out using large FIB currents to enable SEM viewing. The “X” fiducial enables automatic slice and image placement. The resulting image stack will have an isotropic voxel resolution of 4 nm.

Imaging and slicing conditions for FIB-SEM data generation in this workflow are found standard on most instruments. During 3D data collection, the in-column (ICD) and through-the-lens (TLD) detectors are used to form backscattered electron images. The TLD is used for fiducial recognition and automated alignment functions. The ICD provides the final image used in data reconstruction due to its superior ability to image fine, crisp membranes with higher contrast. Milling conditions are 30 keV, 0.79 nA with a 30-µm *z*-depth. Final high resolution images are collected using 2-3 keV, 200-400 pA at 4 mm working distance and 3-µs dwell time.

### Large Volume 3DEM using SBF-SEM

Larger volumes are collected via SBF-SEM, but at lower resolution. In general, SBF-SEM utilizes a microtome equipped with a diamond knife *in situ* to the SEM chamber to physically slice the block rather than ion ablation, as in FIB-SEM (Denk & Horstmann, 2004). Figure 7 illustrates the imaging and slicing geometry, where cutting occurs parallel to the block face and images are collected by scanning the electron beam perpendicularly. In the case of SBF-SEM, the same collection software, Maps, can be used for 2D mapping and serial collection of images after physical slicing. The process is automated and several ROIs may be defined over the block face for collection using the same or different imaging conditions with respect to each other. Again, the process repeats to form an image stack which is later aligned and annotated. Data collection reported here uses an FEI Teneo VolumeScope™ SBF-SEM.

**Figure 7:**
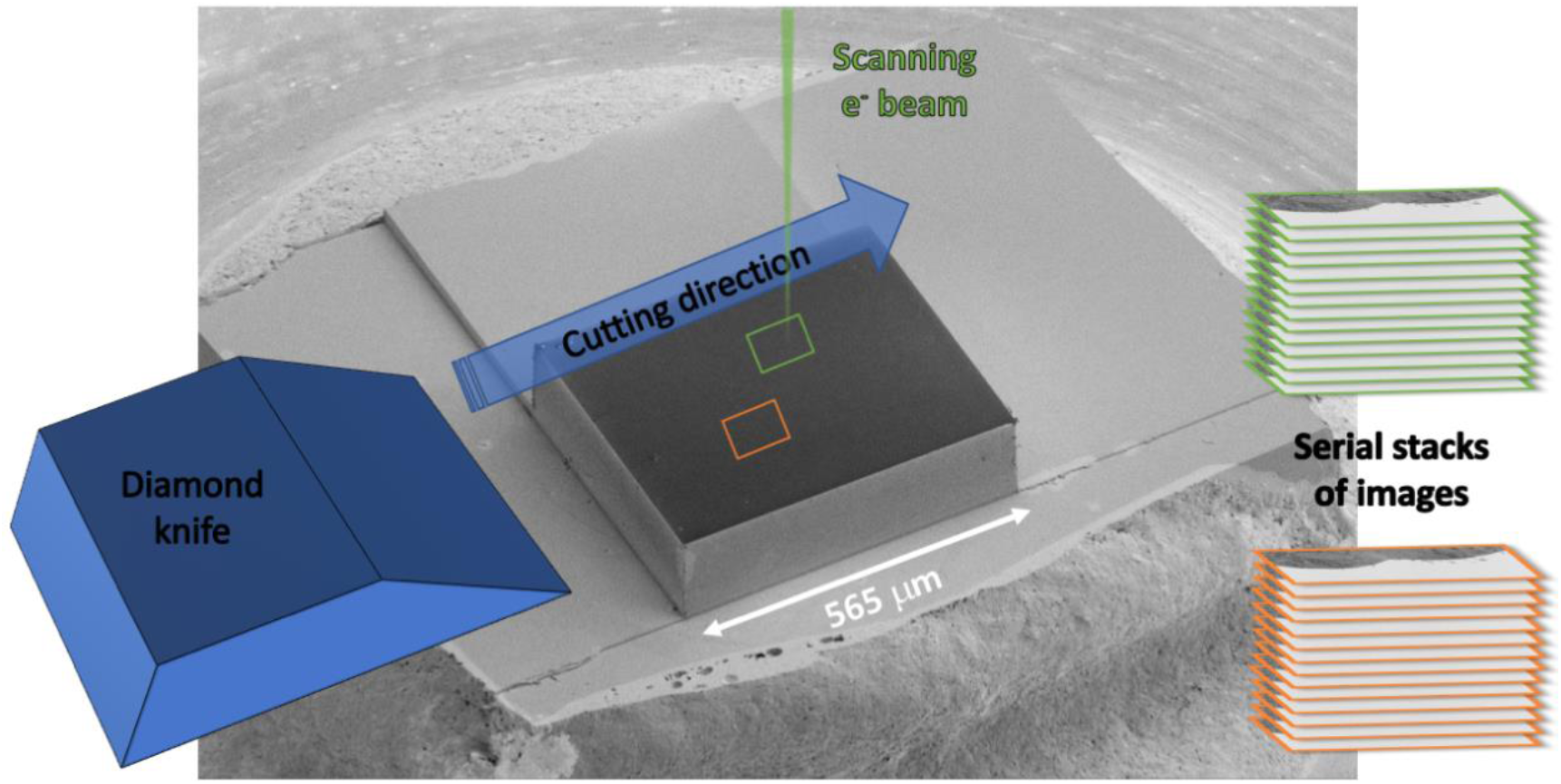
SBF-SEM uses an in situ microtome equipped with a diamond knife to slice across the resin-embedded specimen. Sections fall off the knife to the side of the block. The electron beam is rastered perpendicularly over the freshly exposed block face to form a backscattered electron image. The process continues automatically to form an image stack from the serially sliced sample.

Final imaging conditions for SBF-SEM are more variable due to potentially exposed empty resin at the imaging surface, creating surface charge that needs to be mitigated (Nguyen et al., 2018). Conditions vary widely depending on tissue type or if cell cultures are being imaged. While 40-nm thick slices are always attempted, occasionally slices need to be as thick as 60 nm due to knife chatter or charging influence. Also, variable pressure atmosphere may need to be introduced to mitigate charging and improve cutting performance. Pressure can range between 10 and 50 Pa and is created via water vapor being introduced into the chamber. If tissue is dense and well stained, as with brain, standard high vacuum conditions may be used. It is important to note, however, that most tissues do not image well in high vacuum due to open microstructures and tissue heterogeneity. Scanning conditions are typically 1.5-2 keV, 50-200 pA, at 7 mm working distance using the pole-mounted VolumeScope directional backscattered (VS-DBS) electron detector. Final images are collected such that the complete ROI volume is in a single-frame image tileset, where 10-nm per pixel is the highest achievable spatial resolution. Dwell time ranges between 500 ns and 3-µs and the researcher may utilize several frames of image averaging to improve the signal-to-noise ratio.

### Data Analysis

Currently, 3DEM and large area maps generated are qualitatively analyzed. Data are segmented manually using two software packages: 1) Thermo Scientific Amira (license commercially available) (Pruggnaller, Mayr, & Frangakis, 2008) and 2) Microscopy Image Browser freeware (Belevich, Joensuu, Kumar, Vihinen, & Jokitalo, 2016). Many more options exist and the reader is referred to the paper by Borrett and Hughes, which also describes a common 3DEM processing techniques (Borrett & Hughes, 2016). Amira and code developed in-house using a stochastic version of TurboReg are used to register individual slices with respect to each other for stack alignment (Thévenaz, 2011). Manual segmentation of 3D data sets is time consuming, tedious, and does not keep up with the needed turn-around time to meaningfully inform clinical decisions. Ideally, the process would be automated via machine learning methods. While great strides have been made in the area of connectomics (Boergens et al., 2017; Liu et al., 2014), the same computer vision algorithms do not easily adapt to non-neural tissue. Recently reported methods to minimize segmentation via convolutional neural networks (CNN) are promising (Falk et al., 2018). In-house methods to automatically predict nuclei and nucleoli were utilized on data presented here and are currently in preparation for publication (Loftis et al., 2019). Automated segmentation of large data requires ground truth for training, and manual segmentation is still necessary on a handful of images before prediction algorithms can be applied. Future work will attempt crowd sourcing for improved time-to-data with respect to manual segmentation (Prestopnik, Crowston, & Wang, 2017). In addition, managed workforces, such as CloudFactory, could be utilized to produce training sets faster with minimal resources (HiveMind & CloudFactory; “The wisdom of crowds, for a fee,” 2018).

Once high-throughput methods are developed, large format maps and 3DEM can be quantified to glean meaningful information that will provide insight to clinical outcome and potential therapies. Cellular and nuclear volumes, organelle density, location, and organization, percentage of heterogeneous tissue, etc. are just a few examples that will be quantified and used to inform clinicians.

## Conclusion

A straightforward workflow making large format 2D mapping and 3DEM routine has been presented. Everything from sample fixation to final data quantification has been outlined as a simple approach. Wherever possible, a summary of limitations was provided and alternative solutions given. The workflow was developed with cancer tissue in mind, but can be extended to cell culture, organoids, and other tissue types from various species. Researchers are encouraged to adapt the protocols outlined here in order to fine tune final image quality.

We have successfully implemented this workflow as a research assay in the clinical trial setting as part of the Serial Measurements of Molecular and Architectural Responses to Therapy (SMMART) program (Johnson et al., 2018) being conducted at OHSU and in related research programs, such as the Brenden-Colson Center for Pancreatic Cancer and the Cancer Systems Biology Consortium. The SMMART program couples 3DEM imaging with various protein analyses, cyclic Immunofluorescence and multiplex IHC imaging on the same biopsy from adjacent sections of tumor cores. Results from these assays are already being compared to one another and theories of therapeutic resistance are being tested via patient-derived cell cultures, organoids, and xenografts. In the future, 3DEM will be further used to understand a tumor’s dynamic behaviors and interactions at a nanometer length scale, with certification as a diagnostic imaging tool being the goal.

## Acknowledgements

All the data collected and workflow development included in this manuscript was performed at the Multiscale Microscopy Core (MMC), an OHSU University Shared Resource with technical support from the OHSU Center for Spatial Systems Biomedicine (OCSSB). The SMMART clinical coordination team was invaluable for specimen acquisition support. The authors would like to thank Dr. Bram Koster from the University of Leiden for fruitful discussions and encouragement. Mrs. Sharon Frase and Ms. Julie Justice of St. Jude Children’s Hospital in Memphis, TN for sharing and discussing protocols. We would also like to thank Ms. Emily Benson and Dr. Grahame Kidd from Renovo Neural, Inc. and Dr. Danielle Jorgens from UC-Berkley for past advice. This manuscript was supported with funding from the Prospect Creek Foundation, Brenden-Colson Center for Pancreatic Care, the NCI Cancer Systems Biology Measuring, Modeling, and Controlling Heterogeneity (M2CH) Center awarded to Joe Gray (5U54CA209988), the NCI Human Tumor Atlas Network (HTAN) Omic and Multidimensional Spatial (OMS) Atlas Center awarded to Joe Gray (5U2CCA233280), and the OCSSB.

This study was approved by the Oregon Health & Science University protocol #16113 Institutional Review Board (IRB). Participant eligibility was determined by the enrolling physician and an informed consent was obtained prior to all study protocol related procedures.

